# Activation and signaling characteristics of the hydroxy-carboxylic acid 3 receptor identified in human neutrophils through a microfluidic flow cell technique

**DOI:** 10.1101/2024.12.17.628937

**Authors:** Huamei Forsman, Wenyan Li, Neele K. Levin, Roger Karlsson, Anders Karlsson, Claes Dahlgren, Martina Sundqvist

## Abstract

Human neutrophils express numerous G protein-coupled receptors (GPCRs) of importance for immune regulation. Several of the functionally known neutrophil GPCRs, are not part of the human neutrophil proteome. To identify GPCRs not earlier shown to be expressed in human neutrophils, we utilized a microfluidic flow cell technique in combination with subcellular granule fractionation and liquid chromatography-tandem mass spectrometry (LC-MS/MS). This approach added the hydroxy-carboxylic acid 3 receptor (HCA_3_R) to the human neutrophil proteome. The β-oxidation intermediate 3-hydroxy-octanoic acid (3-OH-C8) is the primary endogenous agonist for this receptor which is highly expressed in adipocytes where it has anti-lipolytic effects. Literature describing the role and function of HCA_3_R in human neutrophils is scarce. We now show that 3-OH-C8 as well the synthetic HCA_3_R agonist IBC 293 activate human neutrophils determined as an increase in the intracellular concentration of free calcium ions ([Ca^2+^]_i_) and activation of the NADPH oxidase. However, in contrast to the rise in [Ca^2+^]_i_ which could be triggered in naïve neutrophils, pre-treatment of neutrophils was required for HCA_3_Rs to activate the NADPH oxidase. That is, the HCA_3_R-mediated NADPH oxidase activation occurred only in neutrophils pre-treated with either an actin cytoskeleton disrupter or an allosteric modulator targeting the GPCR termed free fatty acid receptor 2 (FFA2R). Our findings demonstrate that HCA_3_R is not only a new member of the human neutrophil proteome, but also exhibits functional activity with complex signaling pathways when stimulated with endogenous and synthetic HCA_3_R agonists.

## 1. Introduction

Neutrophils, the most abundant leukocytes in human blood are of prime importance for our innate immune defense against invading microorganisms and for a proper initiation and resolution of inflammation [1]. Neutrophils sense molecules that regulate the inflammatory activity in large part through expression of G protein-coupled receptors (GPCRs) [1–4]. These GPCRs are exposed on the cell surface and/or localized in the membrane of mobilizable intracellular granules/vesicles in resting (naïve) peripheral blood neutrophils [5]. In response to different environmental changes, e.g., increased levels of pro-inflammatory cytokines such as tumour necrosis factor (TNF) the granule-/vesicle-localized receptors can be mobilized to the cell surface. Neutrophils in which the granule-/vesicle-stored receptors have been mobilized to the cell surface have an increased receptor repertoire that allow them to sense and respond to an increased number of danger-associated molecular patterns (DAMPs, e.g., ATP that bind the GPCR termed P2Y_2_ receptor [P2Y_2_R]) and pathogen-associated molecular patterns (PAMPs, e.g., bacterial derived formyl peptides that bind the GPCRs termed formyl peptide receptors [FPR1 and FPR2]). In addition, these neutrophils are commonly transferred to a functional state characterized by an enhanced responsiveness to certain stimuli, e.g., an augmented (primed) production of superoxide anions (O_2_ ^-^) generated by the NADPH oxidase in response to the FPR1 prototype agonist fMLF [3, 5–8].

With > 800 members, the GPCRs comprise the largest family of human membrane receptors [9, 10]. Functional studies using selective ligands targeting diverse GPCRs have shown that human neutrophils express several different GPCR family members [3]. Yet, in studies aiming to identify human neutrophil constituents using tool systems such as large-scale protein expression, many proteins (> 1000) have indeed been identified. However, only a handful of these are GPCRs and several GPCRs shown to be functionally expressed in human neutrophils are not part of the in the proteome [11]. The GPCRs that have been shown to be functionally active in neutrophil but remain to be identified at the protein level include for example FFA2R (Uniprot accession number [AC]: O15552), P_2_Y2R (Uniprot AC: P41231), HRH2 (Uniprot AC: P25021), CXCR4 (Uniprot AC: P61073), and ChemR23 (Uniprot AC: Q99788). It is, thus, obvious that the sample preparations technique used to proteolytically digest intact proteins in subcellular organelles to peptides for proteome profiling of neutrophils must be improved [11]. It has been described that a microfluidic flow cell technique with which immobilized membranes, organelles or intact cells are used as substrate for the enzymatic digestion of proteins, which may improve handling, characterization and identification of plasma membrane and organelle expressed proteins that are difficult to process by other techniques that typically involve organic solvents or detergents [12]. More importantly, the flow cell technique allows isolated plasma membranes and organelles to be immobilized, and the stationary phase of membrane expressed proteins formed can then be used for proteolytic digestion of the parts of the proteins that are exposed on the surface of the plasma membrane or on the cytosolic side of organelle membranes.

In an attempt to improve the detection level of neutrophil GPCRs exposed on the plasma membrane and on intracellular organelle membranes, we isolated subcellular fractions using a two-step Percoll gradient as described by Niels Borregaard and coworkers [13, 14]. The isolated specific granules and the membranes in the light membrane fraction enriched in plasma membranes together with secretory vesicles were immobilized on the surface of a microfluidic flow cell chamber. Proteolytic cleavage was then performed on the immobilized plasma and granule membranes and the cleaved peptide fragments were analyzed with liquid chromatography-tandem mass spectrometry (LC-MS/MS; [15]). In addition to GPCRs already described to be expressed in neutrophils, we identified peptide sequences identical to parts of the cytosolic C-terminus of the hydroxy-carboxylic acid 3 receptor (HCA_3_R, also known as GPR109B, HM74, and Niacin receptor 2 [NIACR2]), suggesting that this receptor is expressed in neutrophils at the protein level. To characterize HCA_3_R further, we utilized two selective HCA_3_R agonists, the endogenous β-oxidation intermediate 3-hydroxy-octanoic acid (3-OH-C8; PubChem compound identifier [CID]: 26613) [16] and the synthetic molecule termed IBC 293 (1-(1-Methylethyl)-1H-benzotriazole-5-carboxylic acid; PubChem CID: 2736690) [17] and investigated HCA_3_R mediated human neutrophil functions with focus on NADPH oxidase activity. Our results show that despite that both agonists triggered a rise in the intracellular concentration of free calcium ions ([Ca^2+^]_i_) in neutrophils, they were poor activators of the NADPH oxidase when added to either naïve or TNF-primed neutrophils. Yet, when IBC 293 was added to neutrophils with either a disrupted actin cytoskeleton or pre-treated with an allosteric modulator selective for the GPCR termed free fatty acid receptor 2 (FFA2R), this synthetic HCA_3_R agonist was turned into a potent activator of the superoxide anion (O_2_ ^-^)-generating NADPH oxidase. The latter route for IBC 293-induced NADPH oxidase activity relied on a receptor transactivation of allosterically modulated FFA2Rs and the response was functional selective (biased) towards O_2_ ^-^-production as presence of the allosteric FFA2R modulator lacked effect on the IBC 293 mediated increase in [Ca^2+^]_i_.

## 1. Material and Methods

### 2.1. Chemicals

Cytiva Ficoll-Paque^TM^ Plus Medium was from Fischer Scientific (Gothenburg, Sweden), Dextran T500 was from Pharmacosmos (Holbæk, Denmark) whereas Fura-2-acetoxymethyl ester (AM) was from Invitrogen by Thermo Fischer Scientific (Gothenburg, Sweden). Isoluminol, TNF, *N*-formyl-methionyl-leucyl-phenylalanine (fMLF), latrunculin A, propionic acid (propionate), the allosteric FFA2R modulator Cmp58 ((*S*)-2-(4-chlorophenyl)-3,3-dimethyl-*N*-(5-phenylthiazol-2-yl)butanamide) and adenosine 5′-thriphosphate disodium salt hydrate (ATP) were from Sigma-Aldrich (Merck, Burlington, MA, USA). The FFA2R antagonist CATPB ((*S*)-3-(2-(3-Chlorophenyl)acetamido)-4-(4-(trifluoromethyl) phenyl) butanoic acid) was from Tocris Bioscience (Bristol, United Kingdom), horseradish peroxidase (HRP) and bovine serum albumin fraction V (BSA) were from Roche (Merck, Burlington, MA, USA) and the Gα_q_ inhibitor YM-254890 was from Wako Chemicals Europe (Neuss, Germany). The HCA_3_R agonists IBC 293 (1-(1-Methylethyl)-1*H*-benzotriazole-5-carboxylic acid) and 3-hydroxy-octanoic acid (3-OH-C8) were from Tocris (Bristol, UK) and Cayman Chemical (Michigan, USA), respectively.

All ligands and reagents were dissolved, aliquoted and stored as recommended by the manufacturer. For experiments, Krebs-Ringer-Glucose (KRG) buffer (120 mM NaCl, 4.9 mM KCl, 1.5 mM MgSO_4_, 1.7 mM KH_2_PO_4_, 8.3 mM Na_2_HPO_4_, 10 mM glucose with or without 1 mM CaCl_2_ in dH_2_O, pH 7.3) was used for subsequent dilutions of ligands and reagents

### 2.2. Ethics statement

The buffy coats used in this project came from healthy human blood donors and were obtained from the stem cell- and component laboratory at Sahlgrenska University Hospital (Gothenburg, Sweden). In accordance with the Swedish legislation section code 4§ 3p SFS 2003:460, no ethical approval for this project was needed as the buffy coats were provided anonymously, i.e., they cannot be traced back to any specific donor.

### 2.3. Isolation of human peripheral blood neutrophils and subcellular fractionation

Neutrophils from healthy donors were isolated from buffy coats by dextran sedimentation, Ficoll-Paque gradient centrifugation and hypotonic lysis of remaining erythrocytes as described [18, 19]. The cells used for functional analysis were suspended in KRG with CaCl_2_ and stored on ice until use.

Subcellular fractionation was performed as described [20]. In short, the isolated neutrophils were treated with the serine protease inhibitor diisopropyl fluorophosphate (DFP, 8 µM final concentration), resuspended in relaxation buffer and, disintegrated by nitrogen cavitation (Parr Instruments Co., Moline, IL), and the post nuclear supernatant was fractionated on two layer Percoll gradients (1.05 g/L and 1.12 g/L), and centrifuged at 15,000 x *g* for 45 min in a fixed-angle JA-20 Beckman rotor. Fractions of 1.5 mL were collected by aspiration from the bottom of the centrifuge tube. The localization of subcellular granules was determined by granule marker analysis as described [20]. The membranes/vesicles in the pooled γ and β fractions enriched in plasma membranes/secretory vesicles and specific/gelatinase granules, respectively, were then used as the starting material for the proteolytic cleavage of the immobilized membranes/organelles and LC−MS/MS analysis of the cleaved peptide fragments.

### 2.4. Lipid-based Protein Immobilization methodology

To identify human neutrophil membrane expressed proteins we applied a surface-shaving approach on intact plasma membranes/secretory vesicles and specific/gelatinase granules by using the lipid-based protein immobilization (LPI®) flow cell (Nanoxis Consulting AB, Gothenburg, Sweden, www.nanoxisconsulting.com). Experiments were performed as described [12] using the membrane fractions isolated from disintegrated neutrophils. In brief, the neutrophil membranes/organelles were injected into a LPI Hexalane FlowCell (Patent Application No. WO2006068619), using a pipette to fill the FlowCell channel (channel volume of approximately 30 μL). The membranes/organelles were allowed to bind to the FlowCell surface. Enzymatic digestion of the proteins was performed by injecting trypsin (20 μg/mL) into the FlowCell channels and incubating for 1 h at room temperature. The generated peptides were eluted by injecting 200 μL ammonium bicarbonate buffer (20 mM) into the channels. The eluted peptides were collected at the outlet ports and transferred into 2.0 mL Axygen tubes. The peptide solutions were incubated at room temperature overnight and subsequently frozen at - 20°C until analysis by mass spectrometry (MS).

### 2.5. Identification of proteins expressed in human neutrophils by Nano-scale liquid chromatographic tandem mass spectrometry

Samples (pooled γ and β fractions) were desalted (Pierce peptide desalting spin columns, Thermo Fischer Scientific) according to the manufacturer instructions prior to analysis on a QExactive HF mass spectrometer interfaced with Easy-nLC1200 liquid chromatography system (Thermo Fisher Scientific). Peptides were trapped on an Acclaim Pepmap 100 C18 trap column (100 μm x 2 cm, particle size 5 μm, Thermo Fischer Scientific) and separated on an in-house packed analytical column (75 μm x 35 cm, particle size 3 μm, Reprosil-Pur C18, Dr. Maisch) using a gradient from 5% to 80% acetonitrile in 0.2% formic acid over 90 min at a flow of 300 nL/min. The instrument operated in data-dependent mode where the precursor ion mass spectra were acquired at a resolution of 60 000, m/z range 400-1600. The 10 most intense ions with charge states 2 to 4 were selected for fragmentation using HCD at collision energy settings of 28. The isolation window was set to 1.2 Da and dynamic exclusion to 20 s and 10 ppm. MS2 spectra were recorded at a resolution of 30 000 with maximum injection time set to 110 ms.

### 2.6. Proteomic Data Analysis

The proteomic analysis and protein characterization were performed at the Proteomics Core Facility at Sahlgrenska Academy, University of Gothenburg, Sweden. In brief, the data files were searched for identification using Proteome Discoverer version 2.4 (Thermo Fisher Scientific). The data was matched against *Human* SwissProt database, using Mascot version 2.5.1 (Matrix Science) as a search engine. The precursor mass tolerance was set to 5 ppm and fragment mass tolerance to 30 mmu. Tryptic peptides were accepted with one missed cleavage and methionine oxidation was set as variable modification. FixedValue were used for peptide-spectrum match (PSM) validation.

### 2.7. Measurement of changes in cytosolic calcium

For detection and measurements of increased concentration of intracellular free calcium ions [Ca^2+^]_i_ when neutrophils were activated by diverse ligands, the neutrophils were loaded with Fura-2AM as previously described [21]. The main steps were as follows; i) neutrophils at a density of 2 × 10^7^ cells/mL in KRG without CaCl_2_ supplemented with 0.1% (w/v) BSA were incubated with 2 μM Fura-2AM at room temperature for 30 minutes, thereafter ii) the Fura-2 labelled neutrophils were washed twice prior iii) resuspended at a density of 2 × 10^7^ cells/mL in KRG with CaCl_2_. The neutrophils were then kept on ice in darkness prior measurements of [Ca^2+^]_i_ on a Perkin Elmer fluorescence spectrophotometer (LC50B), with excitation wavelengths of 340 nm and 380 nm, an emission wavelength of 509 nm, and slit widths of 5 nm and 10 nm, respectively. Disposable polystyrene cuvettes (4.2 mL) containing 2.475 mL reaction mixture (KRG with CaCl_2_ and 5 × 10^6^ Fura-2-stained neutrophils) were first incubated at 37°C for ten minutes prior a baseline measurement for 20 seconds followed by addition of an agonist (25 µL). The figures including measurements of [Ca^2+^]_i_ are shown as a ratio of the detected fluorescence intensities (340 nm: 380 nm) and/or as the peak [Ca^2+^]_i_ of this ratio. The bar graph showing peak [Ca^2+^]_i_ values (Fig. 3D) display the peak values subtracted by the baseline level, which is the value measured just prior before stimulation.

### 2.8. Measurement of NADPH oxidase derived superoxide anions (O_2_ ^-^)

The production of neutrophil NADPH oxidase derived O_2_ ^-^ was assessed using an isoluminol - enhanced chemiluminescence (CL) assay as previously described [22]. In brief, the NADPH oxidase activity was measured in a six-channel Biolumat LB 9505 apparatus (Berthold CO., Wildbad, Germany), using 4-mL disposable polypropylene tubes containing a 0.9 mL reaction mixture (KRG with CaCl_2_, 10 µg/mL isoluminol, 4 U/mL HRP and 10^5^ isolated neutrophils). The reaction mixture was always equilibrated for five minutes at 37°C, prior a baseline measurement of 20 seconds followed by addition of an agonist (0.1 mL) and measurement of O_2_ ^-^ production recorded as light emission over time. The light emission/O_2_ ^-^ production is expressed in Mega counts per minute (Mcpm). For some experiments the isolated neutrophils were incubated (primed) with TNF (10 ng/mL at 37°C for 20 minutes) prior addition to the reaction mixture and for some experiments an inhibitor or antagonist was added during the five minutes equilibration phase at 37 °C. For summary analyses of NADPH oxidase derived O_2_ ^-^ mediated by one agonist (Fig. 5B and 6B), the peak agonist-induced O_2_ ^-^ production subtracted by the baseline level, i.e., the peak value measured just prior stimulation with the respective agonist is shown. For summary analyses of NADPH oxidase derived O_2_ ^-^ mediated by two agonists added in sequence (Fig. 4E and 6D) the peak of the second agonist-induced O_2_ ^-^ production subtracted by the baseline level just prior the second stimulation is shown.

### 2.9 Analysis of data

Analysis of data was performed with GraphPad Prism10 for macOS, version 10.2.3 (GraphPad Software, San Diego, CA, USA). Data in all experiment was repeated with isolated neutrophils from a minimum of three different healthy blood donors. For Fig. 3C and 4C, the half maximal effective concentration (EC_50_) of IBC 293 was analyzed with curve fitting achieved by non-linear regression by the sigmoidal dose–response equation (variable-slope). Raw-data values were utilized for all statistical calculations and a paired two-tailed Student’s *t*-test was applied for data comparison of two groups whereas a repeated measures one-way ANOVA followed by either Tukey’s or Sidak’s multiple comparisons test for data comparison of more than two groups. The precise statistical tests used for distinct figures are defined in the figure legends where applicable. Statistically significant differences are indicated by \**p* < 0.05, ** *p* < 0.01 and *** *p* < 0.001, no statistically significant differences (*p* > 0.05) are not denoted.

## 3. Results

### 3.1. Identification of the HCA_3_ receptor (HCA_3_R) on human neutrophils through a microfluidic flow cell technique

A microfluidic flow cell technique [12] was used to identify GPCRs not earlier described to be expressed in human neutrophils. This technique, involving lipid-based protein immobilization (LPI) methodology, has earlier been used successfully to identify proteins expressed in the outer membrane of different bacteria [15, 23, 24]. Membrane vesicles derived from a neutrophil light membrane fractions enriched in plasma membranes/secretory vesicles (the γ fraction) and a granule fraction enriched in gelatinase/specific granules (the β fraction), were allowed to attach to the surfaces of channels in the LPI flowcell. This was followed by enzymatic digestion for the generation of peptides to be analyzed by LC MS/MS. Trypsin was added to the immobilized neutrophil membranes within the flowcell channels, whereby the proteins exposed on the cytosolic side of the granule/vesicle membranes and on the plasma membrane surface, respectively, were digested into peptide fragments. The peptide fragments formed were collected from the flowcell by eluting buffer through the channels. The peptides were acidified to stop the trypsin activity, frozen and later analyzed by LC MS/MS (Figure 1).

**Figure 1.**
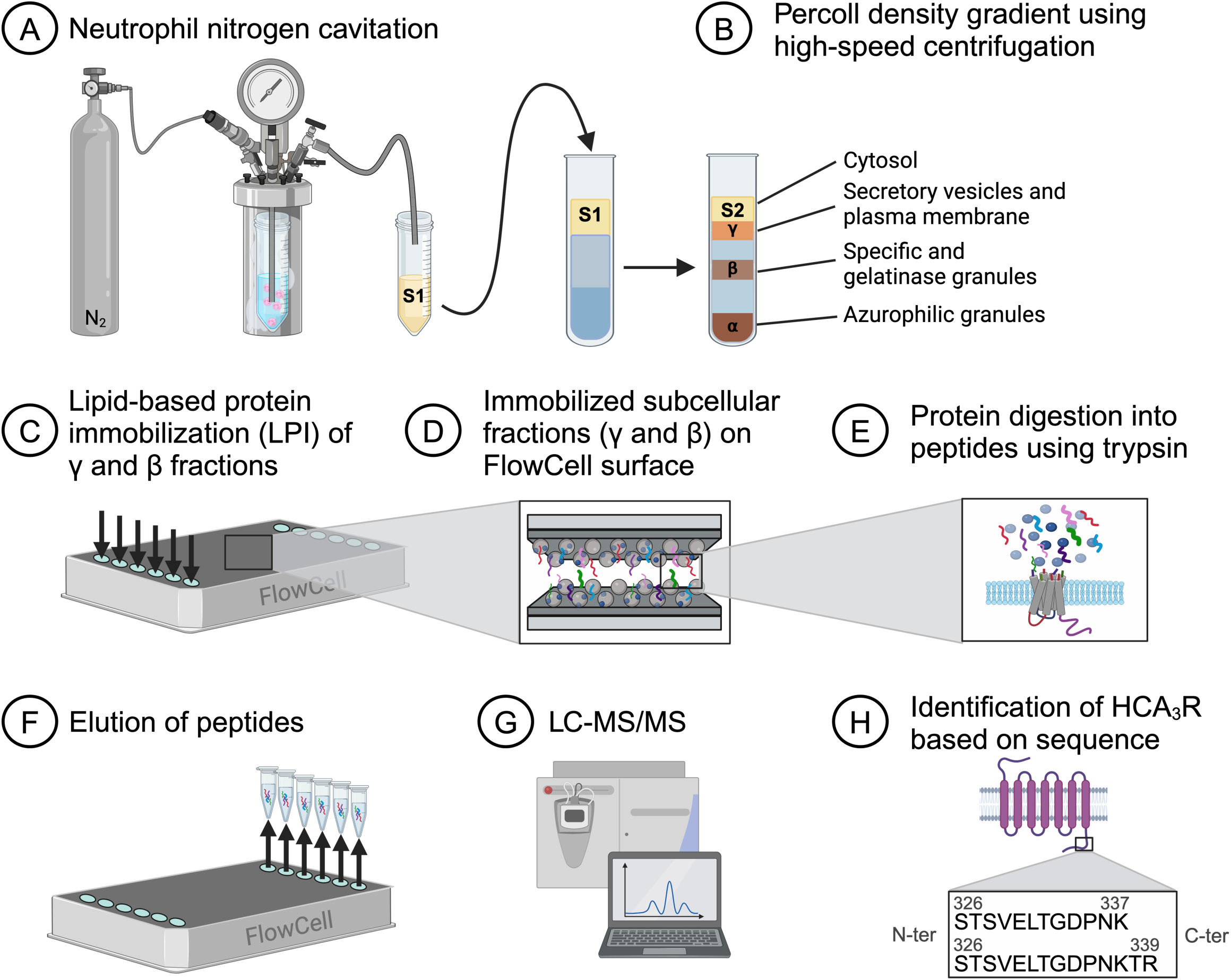
Lipid-based protein immobilization (LPI) methodology and proteomics to identify human neutrophil expressed membrane proteins. A) Isolated human neutrophils were disintegrated by nitrogen cavitation and supernatant 1 (S1) was collected. B) S1 was layered on a Percoll gradient with densities of 1.05 g/mL and 1.12 g/mL and centrifuged to receive the following neutrophil fractions; cytosol (S2), secretory vesicles and plasma membrane (γ), specific and gelatinase granules (β) and azurophil granules (α). C-E) Schematic diagram of the assembly and generation of tryptic peptides in the LPI FlowCell. The samples (γ and β fractions) were allowed to adhere to the surfaces inside the microfluidic flow cell channels as starting material for the immobilized samples prior subjected to proteolytic cleavage (protein digestion into peptides) using trypsin. Two plastic substrates were bonded together with a thin tape spacer that sets the channel height (50 μm). The tape spacer also sets the boundary of the six channels. The samples were injected into the channels of the LPI FlowCell and after attaching to the channels, a trypsin solution was injected into the channels whereof the proteins, including the exposed parts of membrane proteins, were digested for one hour to generate peptide fragments. F) Thereafter a buffer was injected into the channel, whereby the generated peptides were eluted and collected in the outlet port of the channel. G) The cleaved peptide fragments were analyzed with liquid chromatography-tandem mass spectrometry (LC-MS/MS) and H) proteomic analysis revealed two specific peptide sequences that were identical to the amino acid sequence present in the cytosolic tail of HCA_3_R (STSVELTGDPNK and STSVELTGDPNKTR)

Using this LPI methodology, we identified seven GPCRs, previously shown to be expressed both at the protein and functional level in human neutrophils, namely FPR1, FPR2, C5aR, PAFR, BLT_1_R, GPR84 and CXCR2 (Table 1 and [11]). To the best of our knowledge, we also for the first time identified the expression of FFA2R at the protein level in human neutrophils (Table 1). This GPCR has previously only been demonstrated to be functionally expressed in human neutrophils [3, 6]. Moreover, and perhaps more importantly, we identified two specific peptide sequences that were identical to the amino acid sequence present in the cytosolic tail of HCA_3_R (STSVELTGDPNK and STSVELTGDPNKTR; Fig. 1 and Table 1). Similar to FFA2R, neither HCA_3_R has, to the best of our knowledge, previously been identified at the protein level in human neutrophils. Furthermore, the literature regarding if neutrophil expressed HCA_3_Rs are functionally active is scarce.

**Table 1.**
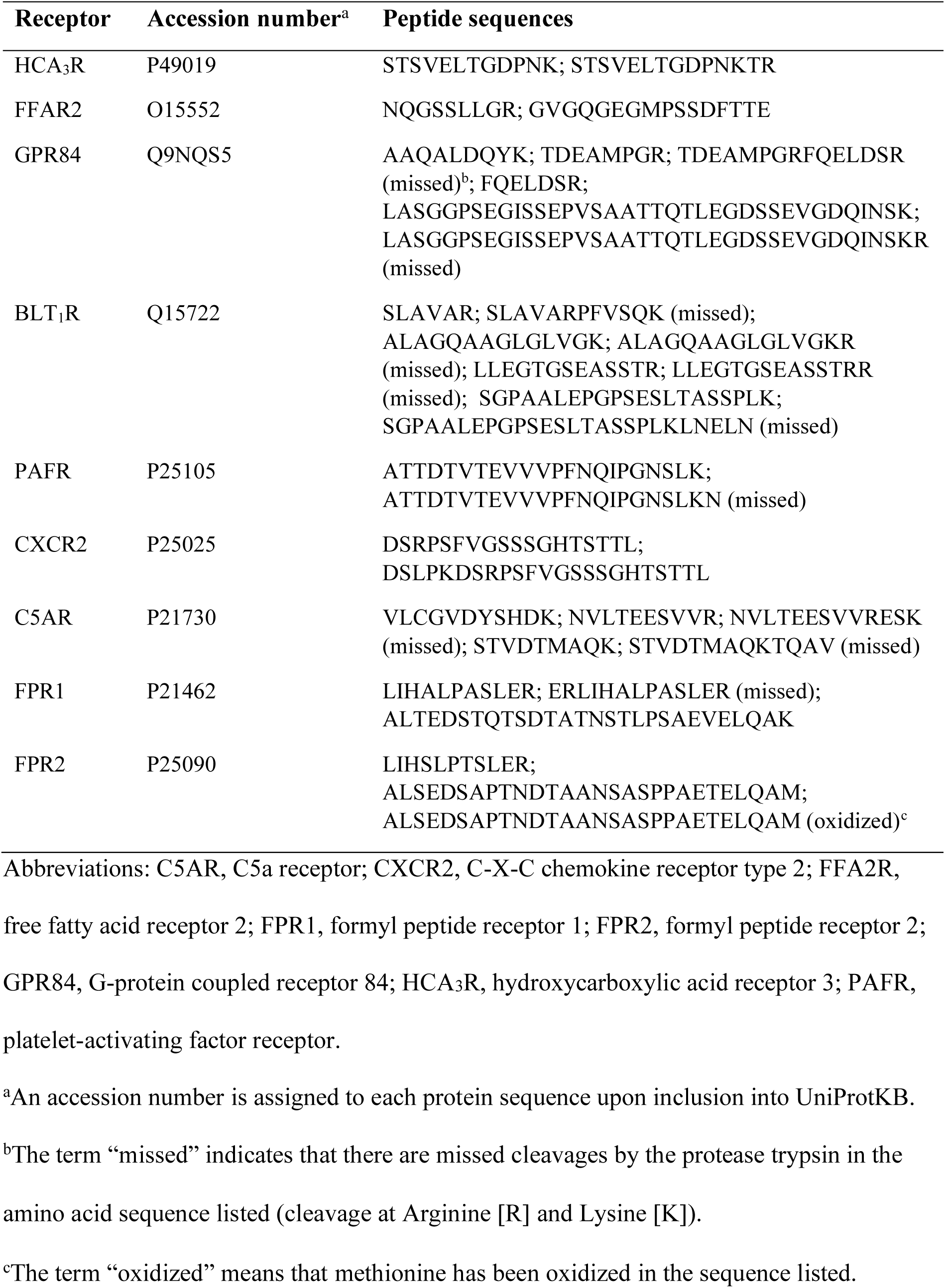
List of GPCRs identified using the microfluidic flow cell technique and their corresponding peptides.

### 3.2. HCA_3_R is functionally expressed in neutrophils – receptor selective agonists mediate a Gα_q_-independent rise in the cytosolic concentration of free Ca^2+^ ([Ca^2+^]_i_)

To investigate the functional activity of HCA_3_Rs expressed in neutrophils, we used 3-hydroxy-octanoate (3-OH-C8), the first known human metabolite to activate HCA_3_Rs in adipocytes [16] along with IBC 293, a synthetic HCA_3_R agonist [17], to activate the cells. The chemical structures of 3-OH-C8 and IBC 293 are shown in Fig. 2. When added to neutrophils, both these HCA_3_R agonists induced a transient increase in [Ca^2+^]_i_. The synthetic agonist IBC 293 was more potent than the endogenous natural agonist 3-OH-C8 (response induced by 5 µM IBC 293 and 600 µM 3-OH-C8, respectively shown in Fig 3A). As no HCA_3_R receptor antagonists have been described that could be used to determine the direct involvement of HCA_3_R in the IBC 293 and 3-OH-C8 mediated increase in [Ca^2+^]_i_, we next performed receptor homologous desensitization experiments. Our data show that neutrophils first activated by IBC 293 were completely desensitized (non-responsive) to a second stimulation with 3-OH-C8 (Fig. 3B). This result strongly implies that both agonists activate the same functionally expressed receptor in human neutrophils, namely HCA_3_R.

**Figure 2.**
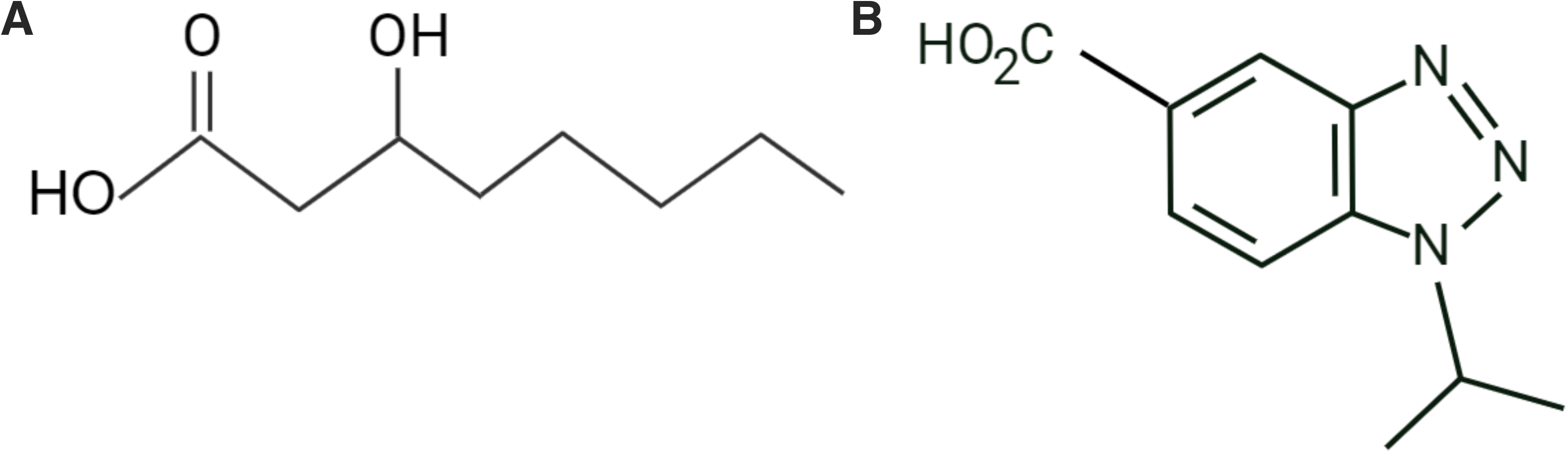
Chemical structures of 3-hydroxy-octanoate (3-OH-C8) and IBC 293. Chemical structures of **A**) the endogenous HCA_3_R agonist 3-hydroxy-octanoate (3-OH-C8) and **B)** the synthetic HCA_3_R agonist IBC 293 (1-(1-Methylethyl)-1H-benzotriazole-5-carboxylic acid).

**Figure 3.**
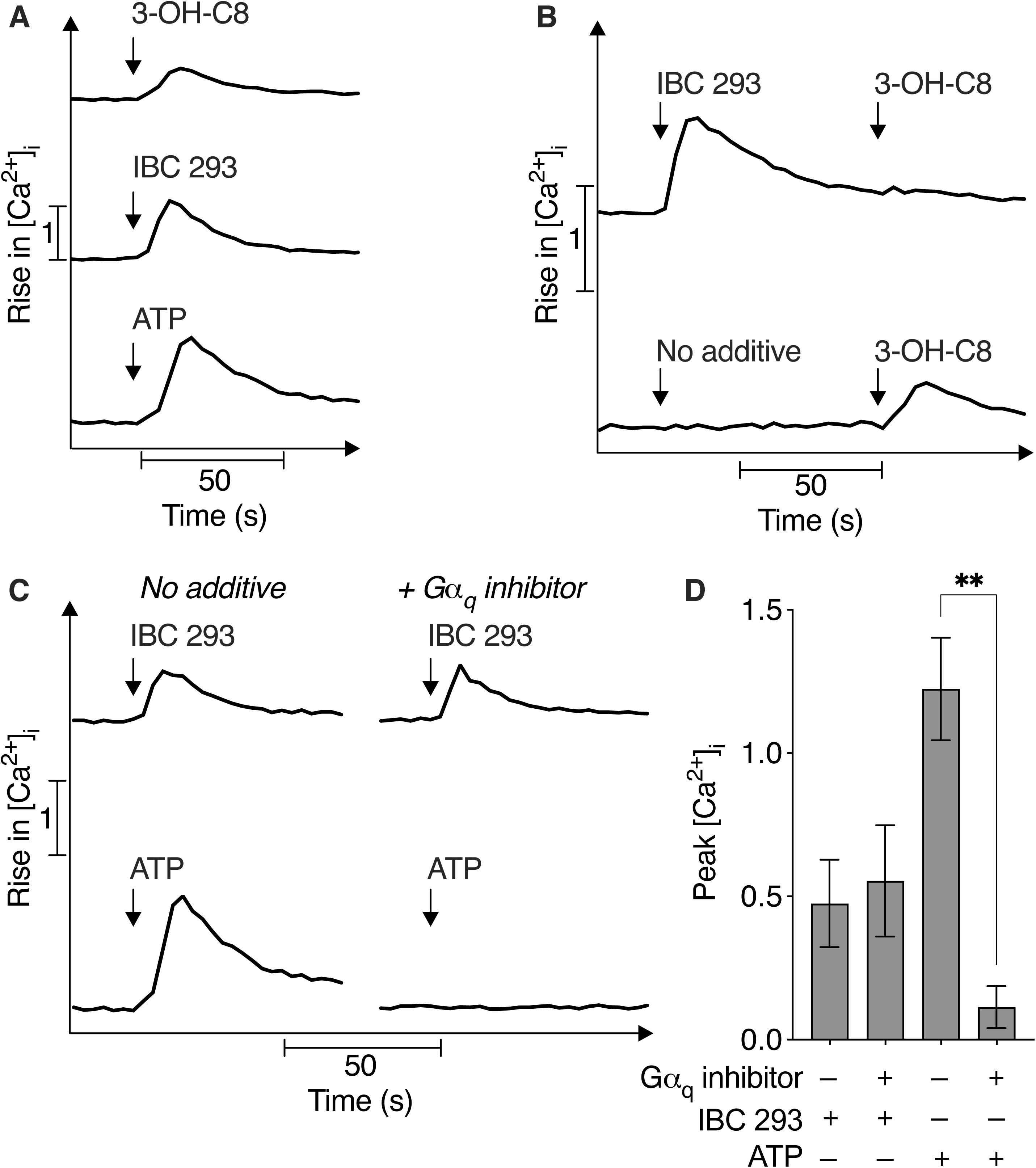
Neutrophils triggered with HCA_3_R selective agonists display a transient rise in intracellular Ca^2+^. Fura 2 loaded neutrophils were pre-incubated at 37°C for ten minutes before stimulated with the HCA_3_R agonists 3-hydroxy-octanoate (3-OH-C8) or IBC 293 or the P2Y_2_R agonist ATP and measurement of a rise in the free intracellular calcium ion (Ca^2+^) concentration [Ca^2+^]_i_ over time. **A**) Neutrophils were stimulated with 3-OH-C8 (600 µM), IBC 293 (5 µM) or ATP (10 µM). One representative trace out of 3-6 individual experiments for each agonist is shown. **B**) Neutrophils were first either stimulated with buffer (no additive) or stimulated with IBC 293 (5 µM). Thereafter, once the IBC 293 mediated rise in [Ca^2+^]_i_ had returned to basal levels, the neutrophils were stimulated a second time with 3-OH-C8 (600 µM). One representative trace out of 3-5 individual experiments for each agonist combination is shown. **C**) Neutrophils were pre-treated (10 min at 37°C) in the absence (no additive; left) or presence of the Gα_q_ inhibitor YM-254890 (200 nM; right) prior stimulation with IBC 293 (2 µM) or ATP (10 µM). **D**) Summary of C showing a comparison of the peak [Ca^2+^]_i_ mediated by IBC 293 (2 µM) or ATP (10 µM) in the absence or presence of 200 nM YM-254890 (mean ± SEM, n = 3). Statistically significant differences were evaluated by a repeated measures one-way ANOVA with Sidak’s multiple comparisons test for analysis of the IBC 293 and ATP mediated peak [Ca^2+^]_i_ in the absence or presence of YM-254890, separately (\*\**p* < 0.01). In A-C, time of agonist addition is indicated by an arrow.

Most neutrophil GPCRs that activate the phospholipase C (PLC) dependent pathway that gives rise to increased [Ca^2+^]_i_ are coupled to a Gα_i_ containing G protein, and PLC is activated by the G protein derived βγ subunit [3]. Two neutrophil receptors that couple to a Gα_q_ containing G protein (the PAF receptor, PAFR, and the ATP receptor, P2Y_2_R; [21, 25, 26]) have, however, been shown to activate the same signaling pathway in which the G protein derived α_q_ subunit activates PLC [3]. To investigate if neutrophil expressed HCA_3_R engage Gα_q_ containing heterotrimeric G proteins for downstream signaling, we used the specific Gα_q_ inhibitor YM-254890 [27]. Pre-treatment of neutrophils with YM-254890 had no effect on the IBC 293 mediated rise in [Ca^2+^]_i_. Activation of neutrophils by the Gα_q_-coupled P2Y_2_R agonist ATP was performed in parallel as a positive control. As expected, presence of the Gα_q_ inhibitor YM-254890 abolished the ATP mediated rise of [Ca^2+^]_i_ (Fig. 3C-D). In summary, these data show that activation of human neutrophil expressed HCA_3_R induces a Gα_q_-independent increase in [Ca^2+^]_i_.

### 3.3. The HCA_3_R agonist IBC 293 activates the neutrophil NADPH oxidase activity through a signaling pathway that is normally blocked by the actin cytoskeleton

It is widely recognized that one of the primary functions of neutrophils is the production of NADPH oxidase derived superoxide anions (O_2_ ^-^; the precursor of other reactive oxygen species [ROS]). These ROS play a crucial role for our host defense through eradication of pathogens but may, if the production is not properly regulated, also harm to surrounding tissues [6, 28]. Our results show that when it comes to untreated isolated human blood neutrophils (naïve neutrophils), the HCA_3_R agonist IBC 293 was a very poor activator of the NADPH oxidase (Fig. 4A). This is also true for the P2Y_2_R agonist ATP, which has been demonstrated as a biased signaling agonist. That is, when ATP binds neutrophil expressed P2Y_2_Rs an activation of the PLC pathway is initiated that leads to a rise in [Ca^2+^]_i_. Yet, this increase in [Ca^2+^]_i_ is not accompanied by activation of the O_2_ ^-^ producing NADPH oxidase. This biased signaling by ATP is not due to an inability of P2Y_2_R to activate the NADPH oxidase but to that this signaling pathway in naïve neutrophils is blocked by the actin cytoskeleton [25]. Accordingly, the NADPH oxidase was activated by ATP when the cytoskeleton first had been disrupted by the actin cytoskeleton disrupting agent latrunculin A (Fig. 4B). Also, IBC 293 triggered a robust O_2_ ^-^ production in neutrophils pre-treated with latrunculin A (Fig. 4A) and the response was concentration dependent with a half maximal effective concentration (EC_50_) of around 9 µM (Fig. 4C) for this synthetic HCA_3_R agonist. However, despite presence of latrunculin A hardly any O_2_^-^ was produced in response to the HCA_3_R agonist 3-OH-C8 (Fig. 4B).

**Figure 4.**
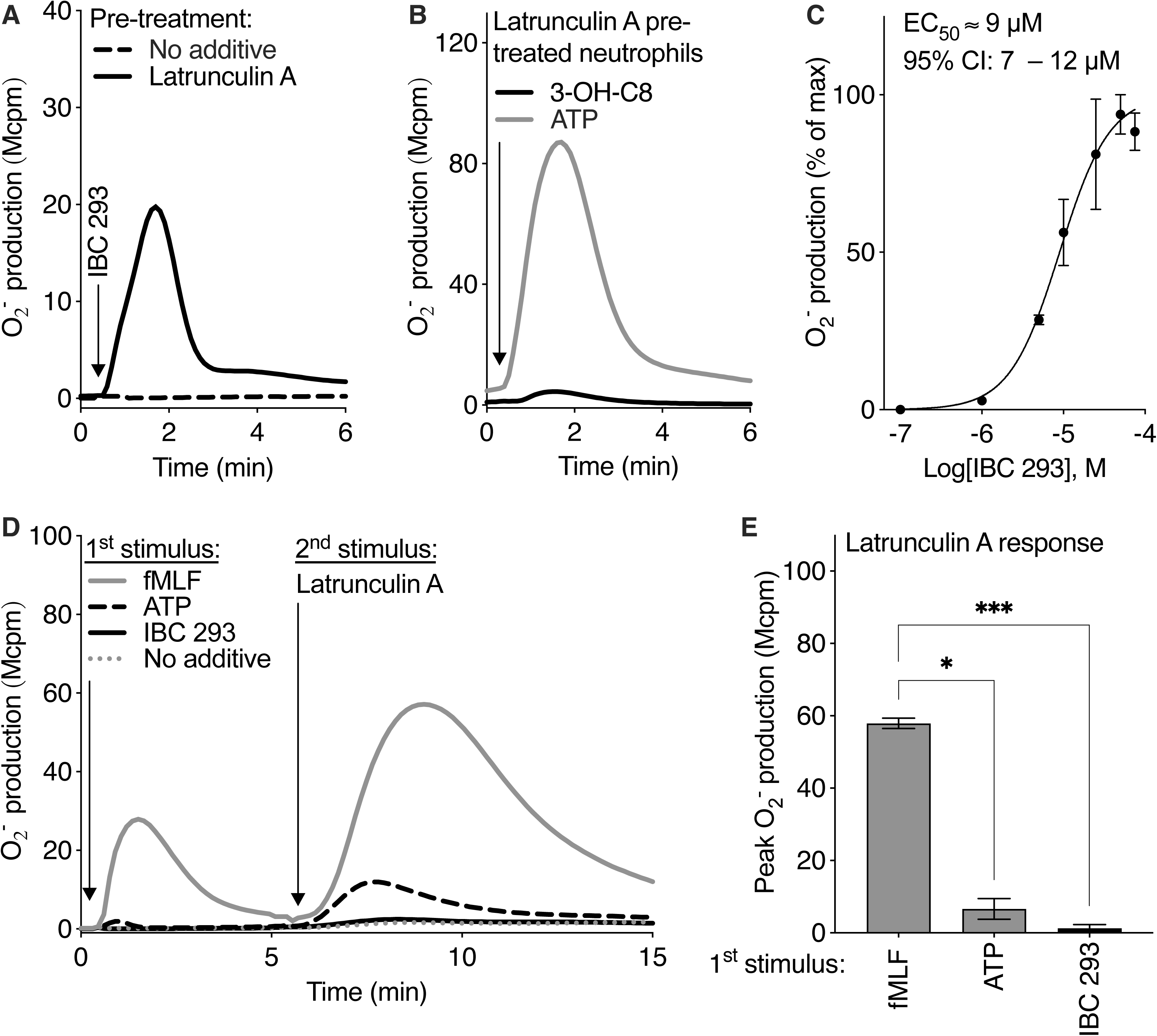
IBC 293 can activate NADPH oxidase provided that the neutrophils have been pre-treated with latrunculin A to disrupt the actin cytoskeleton. **A**-**C**) Neutrophils were pre-pretreated in the absence (no additive) or presence of latrunculin A (25 ng/mL, 37°C, five minutes) prior stimulation the HCA_3_R agonists **A**) IBC 293 (50 µM) **B**) 3-OH-C8 (600 µM), or the P2Y_2_R agonist ATP (50 µM) and measurement of NADPH oxidase produced oxygen radicals (O_2_^-^) over time. One representative trace out of 3-9 individual experiments for each agonist is shown. **C**) A dose-response curve with the half maximal effective concentration (EC_50_) of the IBC 293-induced O ^-^ production by latrunculin A pre-pretreated neutrophils is shown (95 % confidence interval [CI]; mean ± SEM, n = 3). **D**-**E**) Neutrophils, incubated for five minutes at 37°C, were first stimulated with the FPR1 agonist fMLF (100 nM), ATP (50 µM), IBC 293 (50 µM) or buffer (no additive). After around five minutes, when the fMLF response had returned to basal levels, the neutrophils were stimulated with the actin cytoskeleton disrupter latrunculin A (25 ng/mL). One representative trace of the latrunculin A-induced O ^-^ production out of 3 individual experiments for each agonist is shown. **E**) Summary of D showing a comparison of the peak latrunculin A-induced O ^-^ production (mean ± SEM, n = 3). Statistically significant differences were evaluated by repeated measures one-way ANOVA with Tukey’s multiple comparisons test (\**p* < 0.05, \*\**p* < 0.01). In A, B and D, time of agonist/latrunculin A addition is indicated by an arrow.

In addition to amplifying the neutrophil GPCR-mediated NADPH oxidase activity, the actin cytoskeleton is also important for termination of this response. As an example, we have previously shown that some agonist desensitized GPCRs (e.g., FPR1) but not others (e.g., P2Y_2_R) can be re-activated for an additional O_2_ ^-^ production if triggered with latrunculin A [6, 25]. However, in contrast to the latrunculin A mediated reactivation of fMLF (FPR1 agonist) desensitized neutrophils, IBC 293 desensitized neutrophils could, in a similar manner as ATP desensitized neutrophils, not be re-activated to produce O_2_ ^-^ upon disruption of the actin cytoskeleton (Fig. 4D-E). In summary, these data show that IBC 293 not only triggers a rise in [Ca^2+^]_i_, but also can activate neutrophil expressed HCA_3_Rs to produce NADPH oxidase derived O_2_ ^-^. Yet, the latter response only occurs when the cells are activated in the presence of latrunculin A.

### 3.4. HCA_3_R downstream signals transactivate the allosterically modulated free fatty acid 2 receptor (FFA2R) – a novel mechanism to activate the NADPH oxidase

We have previously shown that an allosteric modulator (Cmp58) specific for FFA2R transfers this receptor to state that is sensitive to be transactivated by signals generated by several other neutrophil GPCRs [6, 29–32]. The FFA2R allosteric modulator, thus, functions as a master regulator that amplifies the NADPH oxidase activity induced by agonists specific for other neutrophil GPCRs such as P2Y_2_R, PAFR, BLT_1_R, FPR1 and FPR2. Furthermore, this receptor transactivation (receptor crosstalk) involving FFA2Rs and a GPCRs unrelated to FFA2R has been shown to be most pronounced in TNF primed neutrophils [6, 29–31]. Based on that the IBC 293-induced NADPH oxidase activity was dependent on disruption of the actin cytoskeleton in a similar manner as the ATP-induced NADPH oxidase activity (Fig. 4A-B and [25]), we next investigated if also activated HCA_3_ receptors could transactivate allosterically modulated FFA2Rs. In the absence of Cmp58 hardly any NADPH oxidase activity was generated when TNF-primed neutrophils were stimulated with ATP, IBC 293 or 3-OH-C8. However, in the presence of Cmp58 all these three agonists could activate the NADPH oxidase to produce O_2_ ^-^ (Fig. 5A). In contrast to the lack of 3-OH-C8 mediated NADPH oxidase activity in neutrophils with a disrupted actin cytoskeleton (Fig. 4A), 3-OH-C8 could activate the NADPH oxidase in the presence of Cmp58. Yet, the magnitude of the 3-OH-C8-induced O_2_^-^-production tended to be lower than that induced by IBC 293 in the presence of Cmp58 (Fig. 5A-B), which response was IBC 293 concentration dependent with an EC_50_ of around 3 µM (Fig. 5C).

**Figure 5.**
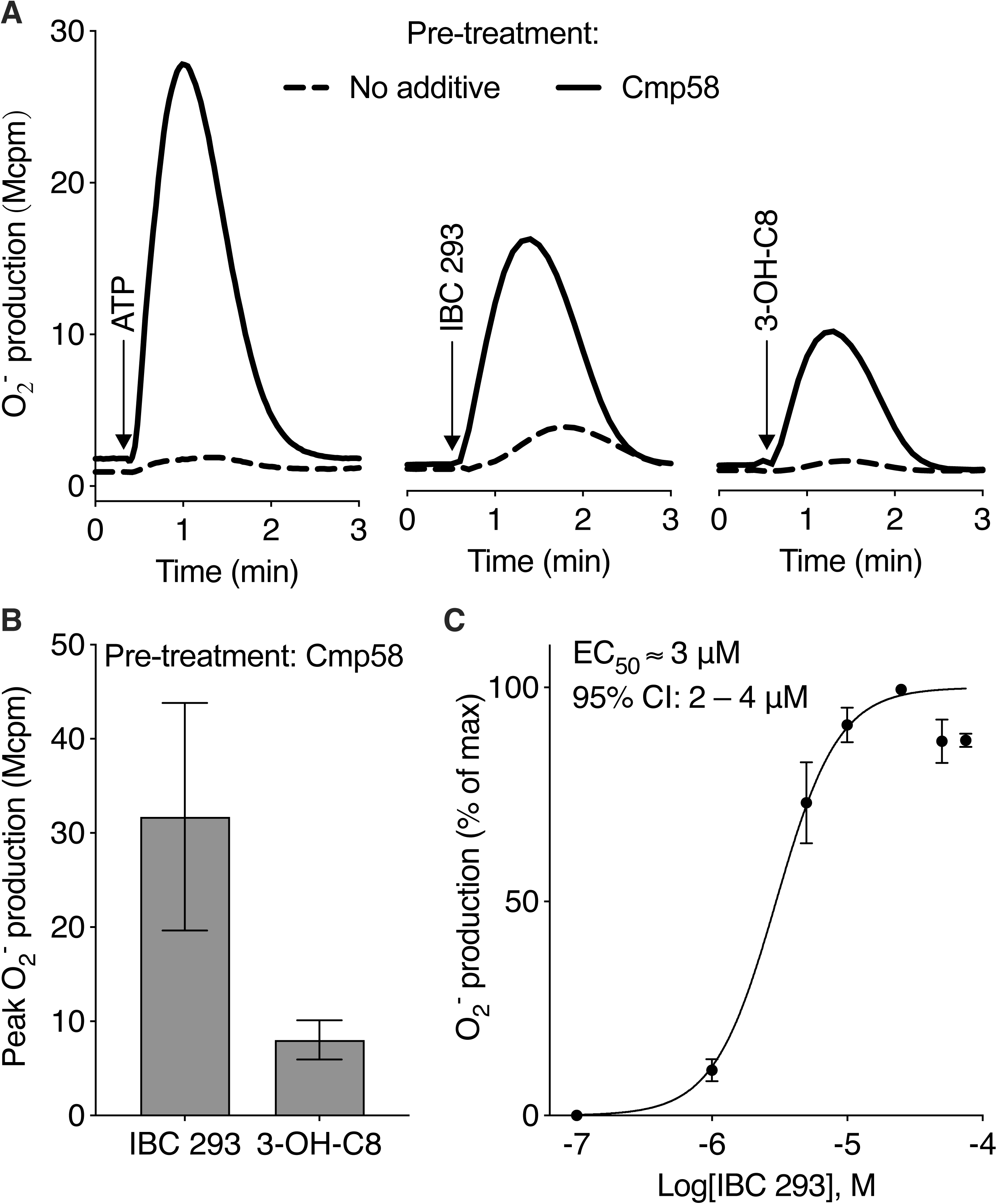
IBC 293 can activate NADPH oxidase in TNF-primed neutrophils with allosterically modulated FFA2Rs. **A**-**C**) Neutrophils were incubated with TNF (10 ng/mL, 37°C, 20 minutes; TNF-primed) and then pre-treated in the absence (no additive) or presence of the FFA2R allosteric modulator Cmp58 (1 µM) at 37°C for five minutes prior stimulation with **A**) the P2Y_2_R agonist ATP (50 µM) or the HCA_3_R agonists IBC 293 (50 µM) or 3-OH-C8 (600 µM) and measurement of NADPH oxidase produced oxygen radicals (O ^-^) over time. One representative trace out of 3-5 individual experiments for each agonist combination is shown. **B**) Summary of A showing a comparison of the peak O ^-^ production induced by IBC 293 or 3-OH-C8 (mean ± SEM, n = 3). **C)** A dose-response curve with the half maximal effective concentration (EC_50_) of the IBC 293-induced O ^-^ production by TNF-primed neutrophils pre-treated with Cmp58 (95 % confidence interval [CI]; mean ± SEM, n = 4). Statistically significant differences in B were evaluated by a paired Student’s *t*-test. In A, the addition of agonist is indicated by an arrow.

To validate the receptor transactivation mechanism between HCA_3_R and FFA2R and to confirm the selectivity of IBC 293 for HCA_3_R but not FFA2R, we utilized the selective FFA2R antagonist CATPB [33, 34]. As expected CATPB inhibited the NADPH oxidase activity induced by the FFA2R orthosteric agonist propionate in the presence of Cmp58 (Fig. 6A-B and [31]), included as a control. In addition, the CATPB also inhibited the IBC 293-induced NADPH oxidase activity in Cmp58 pre-treated neutrophils (Fig. 6A-B). This strongly implies that a receptor transactivation mechanism between HCA_3_Rs and allosterically modulated FFA2Rs triggers down-stream signals that activate the neutrophil NADPH oxidase.

**Figure 6.**
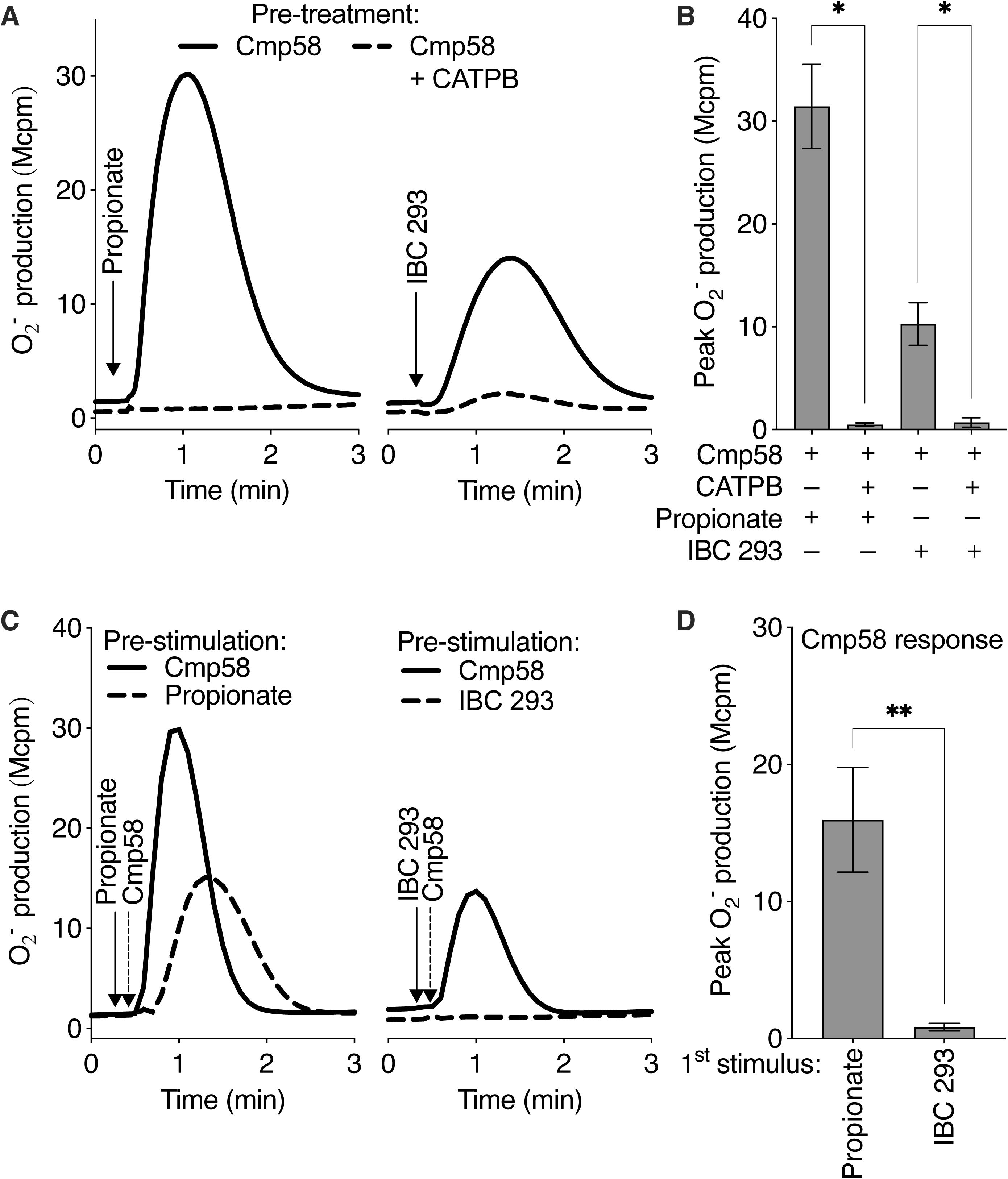
The IBC 293-induced NADPH oxidase activity in the presence of Cmp58 can be inhibited by a selective FFA2R antagonist and is not mediated when the order of Cmp58 and IBC 293 stimulation is reversed. **A**-**B**) Neutrophils were incubated with TNF (10 ng/mL, 37°C, 20 minutes; TNF-primed) and then pre-treated with the FFA2R allosteric modulator Cmp58 (1 µM) in the absence or presence of the FFA2R antagonist CATPB (100 nM) at 37°C for five minutes. Thereafter the neutrophils were stimulated with the FFA2R agonist propionate (25 µM) or the HCA_3_R agonists IBC 293 (25 µM) while measurement of NADPH oxidase produced oxygen radicals (O_2_ ^-^) was recorded over time. One representative trace out of 3 individual experiments for each agonist combination is shown. **B**) Summary data show a comparison of the peak O_2_ ^-^ induced by propionate or IBC 293 production in the in the absence or presence of the CATPB (mean ± SEM, n = 3). **C-D**) TNF-primed neutrophils were first pre-stimulated with Cmp58 (1 µM), the FFA2R agonist propionate (25 µM) or IBC 293 (50 µM). After around five minutes, the neutrophils pre-stimulated with Cmp58 were stimulated a second time with propionate (25 µM) or IBC 293 (50 µM), whereas the neutrophils pre-stimulated with propionate or IBC 293 were stimulated a second time with Cmp58 (1 µM). **C**) One representative out of 6 individual experiments for the O_2_ ^-^ produced after the second stimulation is shown. **D**) Summary of C showing a comparison of the peak O_2_ ^-^ production induced by Cmp58 after the neutrophils had been pre-stimulated (1^st^ stimulus) with propionate or IBC 293 (mean ± SEM, n = 6). Statistically significant differences in B were evaluated by a repeated measures one-way ANOVA with Sidak’s multiple comparisons test for analysis of the propionate and IBC 293 mediated O_2_ ^-^ production in the absence or presence of CATPB, separately and in D by a paired Student’s *t*-test (\**p* < 0.05; \*\**p* < 0.01).) In A and C, the addition of agonist is indicated by an arrow.

To further characterize the mechanism by which the HCA_3_R agonist activates the NADPH oxidase in a Cmp58 dependent manner we investigated the reciprocity of the activation process. The basis for this was that the amplifying effect of a positive allosteric modulator on the response induced by an orthosteric agonist that is recognized by the modulated receptor should, by definition, be reciprocal [29, 35–37]. This is illustrated by the fact that the NADPH oxidase is activated not only when propionate is added to neutrophils with allosterically modulated FFA2Rs but also when the order by which two ligands is added to the neutrophils is revered (Fig 6C-D) and [31, 38]). However, no activation was obtained when the order of stimulation with Cmp58 and IBC 293 was reversed (Fig. 6C-D). This result, together with the desensitization results shown in Fig. 3B demonstrates a selectivity for IBC 293 for neutrophil expressed HCA_3_Rs and implies that the current definition for positive allosteric modulators does not apply to human neutrophils. Taken together, these data show that IBC 293 activated HCA_3_Rs can trigger NADPH oxidase activity through a receptor transactivation mechanism by which the HCA_3_Rs activates allosterically modulated FFA2Rs.

### 3.5. The transactivation between the HCA_3_Rs and FFA2Rs in neutrophils is biased away from a rise in [Ca^2+^]_i_

To further investigate the mechanism behind this receptor transactivation mechanism between HCA_3_R and FFA2R that mediates NADPH oxidase activity we next exploited the capacity of IBC 293 to mediate a transient rise in [Ca^2+^]_i_ in naïve neutrophils (Fig. 3). Our previous data have demonstrated that 250 µM propionate can mediate increased [Ca^2+^]_i_ in the absence of Cmp58. However, we have also shown that presence of Cmp58 can turn low (suboptimal) concentrations of propionate (25 µM), that hardly mediate any increased [Ca^2+^]_i_ into strong mediators of a rise in [Ca^2+^]_i_ (Fig. 7 and [31]). However, similar to what we previously have shown regarding the ATP mediated increase in [Ca^2+^]_i_ [31], presence of Cmp58 had no effect on the IBC 293 mediated rise in [Ca^2+^]_i_. This was regardless of if the neutrophils were stimulated with optimal (5 µM) or suboptimal (0.5 µM) IBC 293 concentrations (Fig. 7). Taken together, these data demonstrate that the signal transduction subsequent the HCA_3_R provoked transactivation of allosterically modulated FFA2R that leads to NADPH oxidase activation is functionally selective and leaves the HCA_3_R mediated rise in [Ca^2+^]_i_ unaffected.

**Figure 7.**
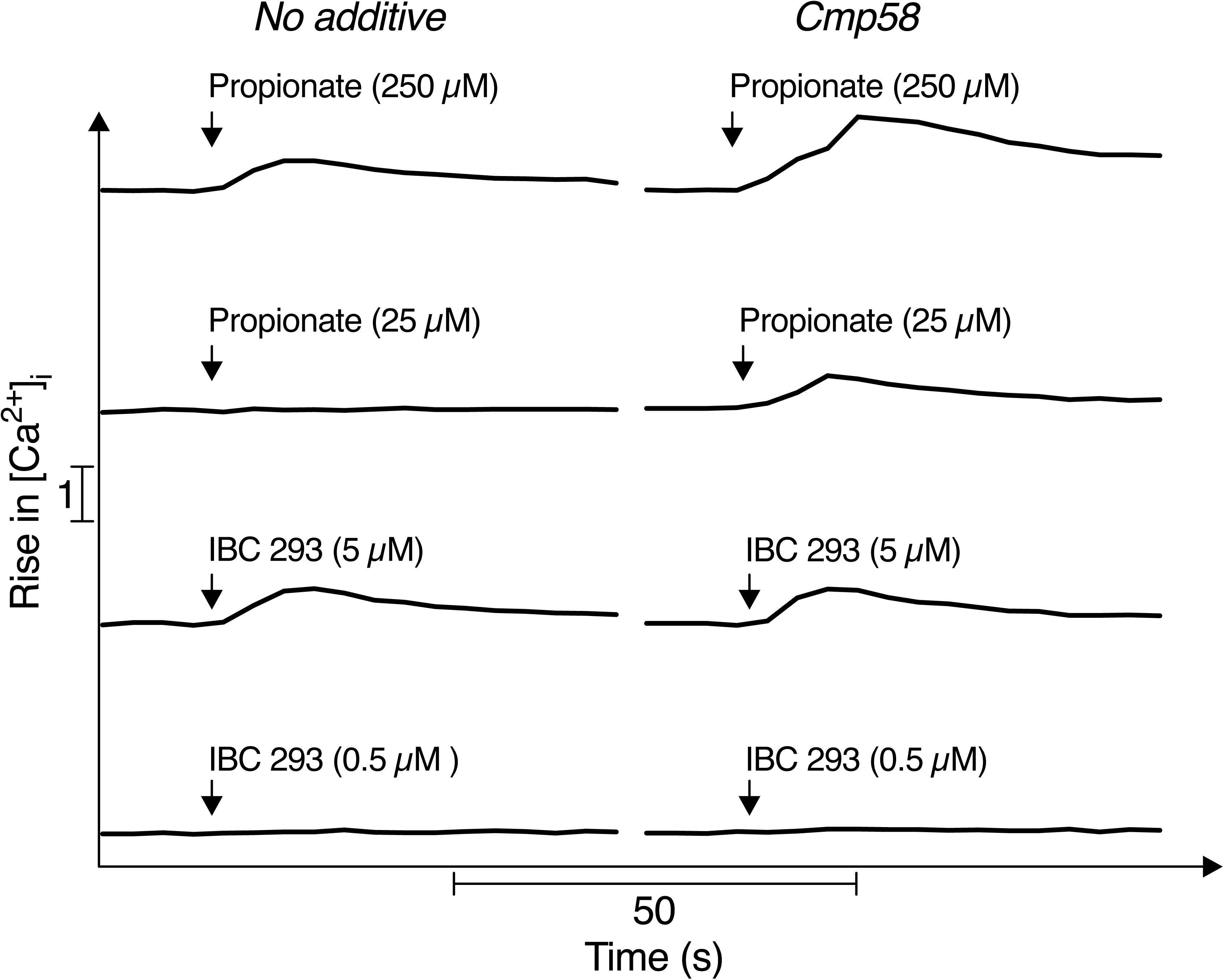
The transactivation mechanism between HCA_3_Rs and FFA2Rs is biased and does not affect the IBC 293 mediated rise in intracellular calcium ions. Fura 2 loaded neutrophils were pre-incubated at 37°C for ten minutes in the absence (no additive) or presence Cmp58 (1 µM). Thereafter the neutrophils were stimulated with propionate (250 µM and 25 µM) or IBC 293 (5 and 0.5 µM) while a rise in the free intracellular calcium ion (Ca^2+^) concentration [Ca^2+^]_i_ was measured over time. One representative out of 3-7 individual experiments for each condition is shown. The addition of agonist is indicated by an arrow.

## 5. Discussion

Neutrophils express many GPCRs, but due to current technical limitations the atlas of human neutrophil GPCRs at the protein level remains incomplete [11]. GPCRs are receptors with multiple transmembrane regions as well as small extracellular regions. This makes it difficult to digest these membrane proteins into peptides suitable for mass spectrometry analysis. In addition, since the expression level of certain GPCRs can be low in some samples, these GPCRs may be underrepresented in samples of high complexity. This means that in some samples and under certain conditions the GPCRs can become “masked” by highly abundant proteins. Therefore, sample preparation strategies are crucial for proteomic analysis. In this study, samples were prepared by subcellular fractionation of human neutrophils, as a “purification” step. These samples were then subjected to the lipid-based protein immobilization (LPI) methodology, which targets membrane proteins in their native state, prior to LC−MS/MS analysis, as an approach to increase the detection sensitivity of GPCRs expressed by human neutrophils. This combination of sample preparation/purification and LPI methodology resulted in the identification of eight GPCRs known to be present and/or functionally expressed in human neutrophils [3, 11]. More importantly, to our knowledge, we also identified the protein expression of HCA_3_R in human neutrophils for the first time. That is, two specific peptide sequences corresponding to the sequence present in the cytosolic tail of HCA_3_R were detected (STSVELTGDPNK and STSVELTGDPNKTR).

HCA_3_R, together with HCA_1_R (GPR81) and HCA_2_R (GPR109A), belongs to a subfamily of GPCRs that are all activated by different energy metabolism intermediates called hydroxycarboxyl acid ligands [39, 40]. The three HCARs share significant sequence homology, but while HCA_1_R binds lactic acid (the end product of glycolysis), HCA_2_R binds the ketone body 3-hydroxy-butyric acid (β-hydroxybutyrate [β-HB]), the HCA_3_R binds the β-oxidation intermediate 3-hydroxy-octanoic acid (3-OH-C8) [16, 41]. All described HCA_3_R-mediated effector functions in other cells are sensitive to pertussis toxin, strongly suggesting that the HCA_3_R couple to Gα_i_/_o_-containing heterotrimeric G proteins [39, 40]. However, while all three HCARs are expressed in adipocytes, only HCA_2_R and HCA_3_R (which share > 95% amino acid sequence similarity) are expressed by immune cells including neutrophils. While there are some studies on the function of neutrophil expressed HCA_2_Rs [39–41], the literature regarding the function of neutrophil expressed HCA_3_Rs is scarce. It has however been implied that neutrophil expressed HCA_3_Rs might be involved in chemotaxis towards certain aromatic D-amino acids (D-phenylalanine and D-tryptophan) [42]. However, the physiological and pathophysiological relevance of this finding has been questioned based on the rare presence of D-amino acids in humans [41]. HCA_3_R was the first of the HCARs whose cDNA was cloned and analyzed. At this time point, almost 30 years ago, it was also reported that neutrophils expressed HCA_3_Rs at the mRNA level [43]. Our current study now adds that neutrophils also express this receptor at the protein level and that it is functionally active in neutrophils when binding its endogenous agonist 3-OH-C8, produced during β-oxidation of fatty acids in various tissues [16, 41]. While the plasma levels measured under healthy conditions appears too low to mediate HCA_3_R activation, the levels of 3-OH-C8 are elevated under conditions of high β-oxidation rates, such as fasting, and it remains to be determined if these elevated levels are sufficient to activate neutrophils in *vivo* [16, 39, 40].

In order to functionally characterize the neutrophil expressed HCA_3_R *in vitro* we used both the endogenous agonist 3-OH-C8 [16], and the synthetic agonist IBC 293 [17]. Apart from determining if these HCA_3_R agonists mediated an increase in [Ca^2+^]_i_, a very early signaling event downstream of activated neutrophil GPCRs, we also investigated agonists ability to activate the O_2_^-^ (the precursor of all reactive oxygen species [ROS]) producing NADPH oxidase. Both 3-OH-C8 and IBC 293 mediated an increase in [Ca^2+^]_i_ when added to naïve neutrophils. However, neither of the two HCA_3_R agonists could activate the neutrophils NADPH oxidase regardless of if the neutrophils were naïve or had ben primed with TNF prior stimulation. This is similar to ATP which upon binding to neutrophil-expressed P2Y_2_Rs mediates a rise in [Ca^2+^]_i_ but lacks capacity to activate the NADPH oxidase in both naïve and primed neutrophils [25]. Hence, our results add to the growing evidence showing that activation of a neutrophil GPCR can mediate increased [Ca^2+^]_i_ without triggering an activation of the NADPH oxidase [6]. That is, a GPCR mediated rise in [Ca^2+^]_i_ is not sufficient for NADPH oxidase activation. Moreover, when it comes to neutrophil expressed GPCRs it appears that the downstream signaling that regulates both the initiation and termination of GPCR-induced NADPH oxidase activity is controlled by the actin cytoskeleton rather than the cytoplasmic adaptor protein β-arrestin [3, 6, 44, 45]. This was evident also in the current study where we show that in a similar manner as ATP also IBC 293 could activate the NADPH oxidase provided that the actin cytoskeleton disrupting agent latrunculin A was present during activation. This implies that the signal transduction downstream of IBC 293 occupied HCA_3_Rs is regulated by the actin cytoskeleton. A pre-requisite for the IBC 293-induced O_2_ ^-^ production was, thus, that the HCA_3_Rs had been uncoupled from the actin cytoskeleton prior to the addition of IBC 293. This suggests that like the downstream signaling of ATP occupied P2Y_2_Rs, the signal transduction downstream of IBC 293 occupied HCA_3_Rs appears to be determined by the actin cytoskeleton. That is, upon actin cytoskeleton disruption, the otherwise blocked signaling route that results in NADPH oxidase activation becomes available. No NADPH oxidase activity was induced when latrunculin A was added to agonist occupied (desensitized) HCA_3_Rs. This lack of re-activation of the agonist occupied HCA_3_Rs by latrunculin A is a character shared with the neutrophil P2Y_2_Rs and the medium chain fatty acid receptor GPR84, but contrasting the effects of latrunculin A on the reactivation of desensitized FPRs [6, 8]. This implies that in contrast to the FPRs, the P2Y_2_Rs, GPR84 and HCA_3_Rs uses a mechanism for termination of the neutrophil NADPH oxidase activity that is independent of the actin cytoskeleton.

The resemblance of the IBC 293 mediated NADPH oxidase activity with that induced by ATP was also evident when neutrophils were stimulated with either of these two agonists in the presence of the FFA2R allosteric modulator Cmp58. We have previously shown that activation of the ATP receptor P2Y_2_R triggers the generation of O_2_ ^-^ only in neutrophils in which their FFA2Rs are allosterically modulated by the presence of Cmp58. The suggested model for this receptor transactivation implies that the signals generated downstream of the activated P2Y_2_R activate the allosterically modulated FFA2Rs from the cytosolic side of the plasma membrane [31]. We now show that a similar receptor transactivation mechanism is used by HCA_3_Rs to activate the NADPH oxidase neutrophils triggered with IBC 293 in the presence of Cmp58. These data not only violate the receptor restriction characteristics generally defining the selectivity of allosteric GPCR modulators but also violate the universal accepted allosteric modulation perception “that a positive (amplifying) as well as a negative (inhibiting) allosteric receptor modulator shall solely affect the response induced by orthosteric agonists specifically recognized by the same receptor as the allosteric modulator” [29, 35–37]. As such the definition for allosteric modulators should, when it comes to neutrophils, be re-evaluated. In addition, the basic function of a positive allosteric modulator also includes that the effect should be reciprocal [29, 35–37]. This means that, with regards to the modulator Cmp58, the amplifying effect should be present irrespective of the sequence in which the allosteric modulator and the orthosteric agonist is added to the cells. This is indeed what characterizes the relationship between Cmp58 and propionate in terms of NADPH oxidase activity (evidenced in this study and [31, 38]). However, in contrast the amplifying effect that Cmp58 has on the IBC 293-induced NADPH oxidase is not reciprocal. This activation pattern in shared with the other neutrophil GPCRs that transactivate the allosterically modulated FFA2Rs and strengthen that the activity of both HCA_3_R and FFA2R are required to obtain an activation of the oxidase. The final proof for the involvement of two different receptors would be to include receptor specific antagonists; this approach has been used in earlier receptor transactivation studies [30], but to our knowledge, no specific HCA_3_R antagonist that can be used is available. However, the inhibitory effect of the specific FFA2R antagonist CATPB (complete inhibition of the transactivation) and the lack of reciprocity of the IBC 293 induced response, together suggest that signals generated both by HCA_3_R and FFA2R are required for the NADPH oxidase activity.

Compared to IBC 293, 3-OH-C8 was a rather poor neutrophil activating agonist and the concentration required for activation was very high. The concentrations of 3-OH-C8 in peripheral blood plasma is normally below 0.4 µM and rarely reach concentrations higher than 10 µM during fasting and other conditions with high β-oxidation rates [16, 46]. The potential physiology/pathophysiology of 3-OH-C8 mediated neutrophil effector functions can, thus, at this stage only be speculated on.

Taken together, we present a surface-shaving approach using a microfluidic flow cell technique enabling identification of GPCRs on human neutrophil membranes. Our study adds also the HCA_3_R as a new member of the human neutrophil proteome and as a functionally active neutrophil-expressed GPCR. Both HCA_3_R agonists tested, the β-oxidation intermediate 3-OH-C8 and the synthetic agonist IBC 293, could mediate increased [Ca^2+^]_i_ and activation of the NADPH oxidase. In conclusion, the results from this study suggest a novel role of neutrophil HCA_3_R-induced ROS production that may be of importance for the regulation of inflammatory processes associated with metabolic changes.

## Glossary

3-OH-C8, HCA3R agonist; CATPB, FFA2R antagonist; Cmp58, FFA2R allosteric modulator; IBC 293, HCA_3_R agonist; YM-254890, Gα_q_ inhibitor; [Ca^2+^]_i_, intracellular concentration of free calcium ions (Ca^2+^)

## Abbreviations

AC: accession number
AM: acetoxymethyl ester
BSA: bovine serum albumin
CID: compound identifier
DAMPs: danger-associated molecular patterns
EC_50_: half maximal effective concentration
FFA2R: free fatty acid receptor 2
fMLF: *N*-formyl-methionyl-leucyl-phenylalanine
FPR: formyl peptide receptor
GPCRs: G protein-coupled receptors
HCAR: hydroxy-carboxylic acid 3 receptor
HRP: horseradish peroxidase
KRG: Krebs-Ringer glucose buffer
LC-MS/MS: liquid chromatography-tandem mass spectrometry
LPI: lipid-based protein immobilization
NIACR2: Niacin receptor 2
O ^−^: superoxide anion
PAMPs: pathogen-associated molecular patterns
PSM: peptide-spectrum match
ROS: reactive oxygen species
TNF: tumour necrosis factor

## CRediT authorship contribution statements

**Huamei Forsman**: Conceptualization, Methodology, Validation, Resources, Investigation, Writing – Review and Editing, Supervision, Project administration, Funding acquisition. **Wenyan Li**: Methodology, Formal analysis, Investigation, Visualization, Writing – Original Draft, Funding acquisition. **Neele Levin**: Methodology, Investigation, Visualization, Writing – Review and Editing. **Roger Karlsson**: Methodology, Validation, Writing – Review and Editing. **Anders Karlsson**: Methodology, Validation, Writing – Review and Editing. **Claes Dahlgren**: Conceptualization, Methodology, Validation, Writing – Review and Editing, Supervision. **Martina Sundqvist**: Conceptualization, Methodology, Validation, Formal analysis, Resources, Investigation, Writing – Original Draft, Visualization, Supervision, Project administration, Funding acquisition.

## Acknowledgments

The work was supported by grants from the Åke Wiberg Foundation (M21-0025, M23-0193 and M24-0227), the Swedish state under the agreement between the Swedish government and the county councils, the ALF-agreement (ALFGBG 78150), the Sahlgrenska International Starting Grant (GU2021/1070), the Swedish Medical Research Council (2018-02848 and 2022-00624), the Swedish Rheumatism Association (R-995669 and R-995361), the Rune and Ulla Almlövs Foundation (2023-418), the Mary von Sydow foundation (2023-4723 and 2024-163), the King Gustaf the V 80-year foundation (FAI-2021-0804, FAI-2022-0873 and SGI-2023-1001), the Magnus Bergwall foundation (2023-875), the Health & Medical Care Committee of the Region Västra Götaland (VGFOUREG-979715 and VGFOUREG-995348), the Wilhelm and Martina Lundgren Science Fund (2024-SA-4605 and 2024-SA-4621), the National Natural Science Foundation of China (82202011) and the Research Foundation of Guangzhou Women and Children’s Medical Center for Clinical Doctors (2020BS003). The sponsors did not have any role in any part of the study. The authors thank Frida Nilsson and Linda Bergqvist for assistance regarding experiments.

## References

[1] A. Mantovani, M.A. Cassatella, C. Costantini, S. Jaillon, Neutrophils in the activation and regulation of innate and adaptive immunity, Nat Rev Immunol, 11 (2011) 519–531.

[2] T. Lämmermann, W. Kastenmüller, Concepts of GPCR-controlled navigation in the immune system, Immunol Rev, 289 (2019) 205–231.

[3] C. Dahlgren, S. Lind, J. Mårtensson, L. Björkman, Y. Wu, M. Sundqvist, H. Forsman, G protein coupled pattern recognition receptors expressed in neutrophils: Recognition, activation/modulation, signaling and receptor regulated functions, Immunol Rev, 314 (2023) 69–92.

[4] H. Forsman, C. Dahlgren, J. Mårtensson, L. Björkman, M. Sundqvist, Function and regulation of GPR84 in human neutrophils, Br J Pharmacol, 181 (2024) 1536–1549.

[5] J.B. Cowland, N. Borregaard, Granulopoiesis and granules of human neutrophils, Immunol Rev, 273 (2016) 11–28.

[6] C. Dahlgren, H. Forsman, M. Sundqvist, L. Björkman, J. Mårtensson, Signaling by Neutrophil G Protein-Coupled Receptors that Regulate the Release of Superoxide Anions, J Leukoc Biol, (2024).

[7] I. Miralda, S.M. Uriarte, K.R. McLeish, Multiple Phenotypic Changes Define Neutrophil Priming, Front Cell Infect Microbiol, 7 (2017) 217.

[8] M. Sundqvist, K. Christenson, A. Holdfeldt, M. Gabl, J. Mårtensson, L. Björkman, R. Dieckmann, C. Dahlgren, H. Forsman, Similarities and differences between the responses induced in human phagocytes through activation of the medium chain fatty acid receptor GPR84 and the short chain fatty acid receptor FFA2R, Biochim Biophys Acta Mol Cell Res, 1865 (2018) 695–708.

[9] S.B. Gacasan, D.L. Baker, A.L. Parrill, G protein-coupled receptors: the evolution of structural insight, AIMS Biophys, 4 (2017) 491–527.

[10] K.L. Pierce, R.T. Premont, R.J. Lefkowitz, Seven-transmembrane receptors, Nat Rev Mol Cell Biol, 3 (2002) 639–650.

[11] S. Rørvig, O. Østergaard, N.H. Heegaard, N. Borregaard, Proteome profiling of human neutrophil granule subsets, secretory vesicles, and cell membrane: correlation with transcriptome profiling of neutrophil precursors, J Leukoc Biol, 94 (2013) 711–721.

[12] E.T. Jansson, C.L. Trkulja, J. Olofsson, M. Millingen, J. Wikström, A. Jesorka, A. Karlsson, R. Karlsson, M. Davidson, O. Orwar, Microfluidic flow cell for sequential digestion of immobilized proteoliposomes, Anal Chem, 84 (2012) 5582–5588.

[13] N. Borregaard, J.M. Heiple, E.R. Simons, R.A. Clark, Subcellular localization of the b-cytochrome component of the human neutrophil microbicidal oxidase: translocation during activation, J Cell Biol, 97 (1983) 52–61.

[14] S.N. Clemmensen, L. Udby, N. Borregaard, Subcellular fractionation of human neutrophils and analysis of subcellular markers, Methods Mol Biol, 1124 (2014) 53–76.

[15] A. Karlsson, L.A. Alarcón, B. Piñeiro-Iglesias, G. Jacobsson, S. Skovbjerg, E.R.B. Moore, P.K. Kopparapu, T. Jin, R. Karlsson, Surface-Shaving of Staphylococcus aureus Strains and Quantitative Proteomic Analysis Reveal Differences in Protein Abundance of the Surfaceome, Microorganisms, 12 (2024).

[16] K. Ahmed, S. Tunaru, C.D. Langhans, J. Hanson, C.W. Michalski, S. Kölker, P.M. Jones, J.G. Okun, S. Offermanns, Deorphanization of GPR109B as a receptor for the beta-oxidation intermediate 3-OH-octanoic acid and its role in the regulation of lipolysis, J Biol Chem, 284 (2009) 21928–21933.

[17] G. Semple, P.J. Skinner, M.C. Cherrier, P.J. Webb, C.R. Sage, S.Y. Tamura, R. Chen, J.G. Richman, D.T. Connolly, 1-Alkyl-benzotriazole-5-carboxylic acids are highly selective agonists of the human orphan G-protein-coupled receptor GPR109b, J Med Chem, 49 (2006) 1227–1230.

[18] A. Böyum, Isolation of mononuclear cells and granulocytes from human blood. Isolation of monuclear cells by one centrifugation, and of granulocytes by combining centrifugation and sedimentation at 1 g, Scand J Clin Lab Invest Suppl, 97 (1968) 77–89.

[19] A. Bøyum, Isolation of lymphocytes, granulocytes and macrophages, Scand J Immunol, Suppl 5 (1976) 9–15.

[20] L. Kjeldsen, H. Sengelov, N. Borregaard, Subcellular fractionation of human neutrophils on Percoll density gradients, J Immunol Methods, 232 (1999) 131–143.

[21] L. Björkman, H. Forsman, L. Bergqvist, C. Dahlgren, M. Sundqvist, Larixol is not an inhibitor of Gα(i) containing G proteins and lacks effect on signaling mediated by human neutrophil expressed formyl peptide receptors, Biochem Pharmacol, 220 (2024) 115995.

[22] C. Dahlgren, H. Björnsdottir, M. Sundqvist, K. Christenson, J. Bylund, Measurement of Respiratory Burst Products, Released or Retained, During Activation of Professional Phagocytes, Methods Mol Biol, 2087 (2020) 301–324.

[23] R. Karlsson, L. Gonzales-Siles, M. Gomila, A. Busquets, F. Salvà-Serra, D. Jaén-Luchoro, H.E. Jakobsson, A. Karlsson, F. Boulund, E. Kristiansson, E.R.B. Moore, Proteotyping bacteria: Characterization, differentiation and identification of pneumococcus and other species within the Mitis Group of the genus Streptococcus by tandem mass spectrometry proteomics, PLoS One, 13 (2018) e0208804.

[24] E. Ahlberg, M.C. Jenmalm, A. Karlsson, R. Karlsson, L. Tingö, Proteome characterization of extracellular vesicles from human milk: Uncovering the surfaceome by a lipid-based protein immobilization technology, J Extracell Biol, 3 (2024) e70020.

[25] M. Gabl, M. Winther, A. Welin, A. Karlsson, T. Oprea, J. Bylund, C. Dahlgren, H. Forsman, P2Y2 receptor signaling in neutrophils is regulated from inside by a novel cytoskeleton-dependent mechanism, Exp Cell Res, 336 (2015) 242–252.

[26] K. Önnheim, K. Christenson, M. Gabl, J.C. Burbiel, C.E. Müller, T.I. Oprea, J. Bylund, C. Dahlgren, H. Forsman, A novel receptor cross-talk between the ATP receptor P2Y2 and formyl peptide receptors reactivates desensitized neutrophils to produce superoxide, Exp Cell Res, 323 (2014) 209–217.

[27] A. Nishimura, K. Kitano, J. Takasaki, M. Taniguchi, N. Mizuno, K. Tago, T. Hakoshima, H. Itoh, Structural basis for the specific inhibition of heterotrimeric Gq protein by a small molecule, Proc Natl Acad Sci U S A, 107 (2010) 13666–13671.

[28] E. Naish, A.J. Wood, A.P. Stewart, M. Routledge, A.C. Morris, E.R. Chilvers, K.M. Lodge, The formation and function of the neutrophil phagosome, Immunol Rev, 314 (2023) 158–180.

[29] C. Dahlgren, A. Holdfeldt, S. Lind, J. Mårtensson, M. Gabl, L. Björkman, M. Sundqvist, H. Forsman, Neutrophil Signaling That Challenges Dogmata of G Protein-Coupled Receptor Regulated Functions, ACS Pharmacol Transl Sci, 3 (2020) 203–220.

[30] S. Lind, K.L. Granberg, H. Forsman, C. Dahlgren, The allosterically modulated FFAR2 is transactivated by signals generated by other neutrophil GPCRs, PLoS One, 18 (2023) e0268363.

[31] S. Lind, A. Holdfeldt, J. Mårtensson, M. Sundqvist, L. Björkman, H. Forsman, C. Dahlgren, Functional selective ATP receptor signaling controlled by the free fatty acid receptor 2 through a novel allosteric modulation mechanism, Faseb j, 33 (2019) 6887–6903.

[32] Y. Wu, C. Dahlgren, H. Forsman, M. Sundqvist, LTB4 is converted into a potent human neutrophil NADPH oxidase activator via a receptor transactivation mechanism in which the BLT1 receptor activates the free fatty acid receptor 2, bioRxiv (2024) 10.1101/2024.1111.1112.623248

[33] L. Björkman, J. Mårtensson, M. Winther, M. Gabl, A. Holdfeldt, M. Uhrbom, J. Bylund, A. Højgaard Hansen, S.K. Pandey, T. Ulven, H. Forsman, C. Dahlgren, The Neutrophil Response Induced by an Agonist for Free Fatty Acid Receptor 2 (GPR43) Is Primed by Tumor Necrosis Factor Alpha and by Receptor Uncoupling from the Cytoskeleton but Attenuated by Tissue Recruitment, Mol Cell Biol, 36 (2016) 2583–2595.

[34] B.D. Hudson, M.E. Due-Hansen, E. Christiansen, A.M. Hansen, A.E. Mackenzie, H. Murdoch, S.K. Pandey, R.J. Ward, R. Marquez, I.G. Tikhonova, T. Ulven, G. Milligan, Defining the molecular basis for the first potent and selective orthosteric agonists of the FFA2 free fatty acid receptor, J Biol Chem, 288 (2013) 17296–17312.

[35] A. Bock, M. Bermudez, Allosteric coupling and biased agonism in G protein-coupled receptors, Febs j, 288 (2021) 2513–2528.

[36] M. Grundmann, E. Bender, J. Schamberger, F. Eitner, Pharmacology of Free Fatty Acid Receptors and Their Allosteric Modulators, Int J Mol Sci, 22 (2021).

[37] D. Wootten, A. Christopoulos, P.M. Sexton, Emerging paradigms in GPCR allostery: implications for drug discovery, Nat Rev Drug Discov, 12 (2013) 630–644.

[38] J. Mårtensson, A. Holdfeldt, M. Sundqvist, M. Gabl, T.P. Kenakin, L. Björkman, H. Forsman, C. Dahlgren, Neutrophil priming that turns natural FFA2R agonists into potent activators of the superoxide generating NADPH-oxidase, J Leukoc Biol, 104 (2018) 1117–1132.

[39] K. Ahmed, S. Tunaru, S. Offermanns, GPR109A, GPR109B and GPR81, a family of hydroxy-carboxylic acid receptors, Trends Pharmacol Sci, 30 (2009) 557–562.

[40] S. Offermanns, Hydroxy-Carboxylic Acid Receptor Actions in Metabolism, Trends Endocrinol Metab, 28 (2017) 227–236.

[41] S. Offermanns, S.L. Colletti, T.W. Lovenberg, G. Semple, A. Wise, I.J. Ap, International Union of Basic and Clinical Pharmacology. LXXXII: Nomenclature and Classification of Hydroxy-carboxylic Acid Receptors (GPR81, GPR109A, and GPR109B), Pharmacol Rev, 63 (2011) 269–290.

[42] Y. Irukayama-Tomobe, H. Tanaka, T. Yokomizo, T. Hashidate-Yoshida, M. Yanagisawa, T. Sakurai, Aromatic D-amino acids act as chemoattractant factors for human leukocytes through a G protein-coupled receptor, GPR109B, Proc Natl Acad Sci U S A, 106 (2009) 3930–3934.

[43] H. Nomura, B.W. Nielsen, K. Matsushima, Molecular cloning of cDNAs encoding a LD78 receptor and putative leukocyte chemotactic peptide receptors, Int Immunol, 5 (1993) 1239–1249.

[44] R.J. Lefkowitz, E.J. Whalen, beta-arrestins: traffic cops of cell signaling, Curr Opin Cell Biol, 16 (2004) 162–168.

[45] L.M. Luttrell, R.J. Lefkowitz, The role of beta-arrestins in the termination and transduction of G-protein-coupled receptor signals, J Cell Sci, 115 (2002) 455–465.

[46] C.G. Costa, L. Dorland, U. Holwerda, I.T. de Almeida, B.T. Poll-The, C. Jakobs, M. Duran, Simultaneous analysis of plasma free fatty acids and their 3-hydroxy analogs in fatty acid beta-oxidation disorders, Clin Chem, 44 (1998) 463–471.

